# massiveGenesetsTest: a web tool to run enrichment analysis

**DOI:** 10.1101/2021.02.15.431228

**Authors:** Luigi Cerulo, Stefano Maria Pagnotta

## Abstract

**Motivation:** Inferring biological phenotypes from genomic data and sample clusters is a routinely task usually performed with Gene-Set Enrichment Analysis (GSEA), a tool that queries gene-profiles. In previous work, we scrutinized the approach based on Mann-Witney for Gene-Sets test. We highlighted the Mann-Witney test-statistics sensitivity to uncover weak signals and the drastic decreasing of time complexity.

**Results:** We propose web implementation of the Gene-sets testing based on the Mann-Witney procedure. The test-procedure has reshaped to decrease the computational expense, now about tens of seconds, even if a large collection of gene-sets queries the same gene-profile. The probabilistic interpretation of the normalized test-statistic has been investigated to a better understanding of the enrichment results. A novel prioritization method across enrichment-scores, gene-set dimensions, and p-values, draws attention to relevant gene-sets. The web tool provides both tabular and graphical enrichment results. A complimentary R function allows integrating the enrichment procedure in a complex context.

**Supplementary information:** Example data and guidelines are included in the supporting material of the web-site.

## 1 Introduction

Gene-sets enrichment analysis is a routinely used technique to uncover phenotypes of a gene profile usually associated with the differential expression between two conditions [7]. The inputs of the enrichment analysis are a gene-profile and a gene-set. The gene-profile is a ranked list of genes, usually correlated to the differential level of expression between two sets of samples, such as treatment and control. A gene-set is a collection of genes cooperating to a specific phenotype derived from annotated databases, such as Gene Ontology [1]. From the work of [12], the statistical framework of enrichment analysis is just a significance test where a p-value is assigned to a set of genes considered as a unit: if the *p*-value is below the significance level of the test then the gene-set is associated with the treatment group, assuming the alternative hypothesis to upper tail. The enrichment score is a metric that measures how relevant is the association between the subsets of gene-sets and the treatment group.

GSEA [9] is the most used gene-set enrichment methodology adopting a modified version of the two-sample Kolmogorov-Smirnov test. Its main drawback is the heavy computational load that for *N* gene-sets is O(KN), with K high due the resampling strategy to compute the null distribution. Other methodologies, such as [6] and [12], concerning Mann-Whitney test [11], known as Mann-Witney-Wilcoxon (MWW) or rank-sum test as well, captured our attention for two main reasons: 1) the computational time can be reduced, 2) insufficient attention has been paid to the test-statics which is crucial to interpret the results. We explored this second point in [3] where we proposed a methodology named MWW-GST (Gene Set Test), based on the normalization of the MWW test-statistic defined as Normalized Enrichment Score (NES), that is an estimate of a probability.

In this note we propose an online implementation of MWW-GST in javascript that allowed us to implement a very fast enrichment analysis tool. Essentially, given a gene-profile, and a collection of gene-sets, to speed up the MWW-GST we precompute the rank of the gene-profile, and then we compute the MWW-statistic for each gene-set. As implemented in javascript, the tool does not run on server side, so the speed of the analysis reflects the speed limit of the client web-browser.

## 2. Methodological details

### 2.1 The Normalized Enrichment Score and the *p*-value

The Normalized Enrichment Score (NES) and the *p*-value come from the Mann-Whitney (MW) test [4]. The null hypothesis of this test states that there’s no mutual dominance of the distribution functions, *F_in_*(*x*) and *F_out_*(*x*), that describe the intensities of the genes, respectively in and out the gene-set. The alternative hypothesis states that the distribution function *F_out_*(*x*) dominates *F_in_*(*x*), i.e. 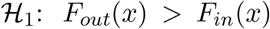. Under the alternative hypothesis, the genes in the gene-set have intensities higher than those of the genes outside the gene-set. The test statistic *U* of the MW-test is the number of times that the relation 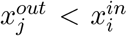 is true ∀ *i,j*, where 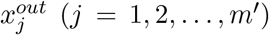 is the intensity associated with the *j^th^* gene outside the geneset, while 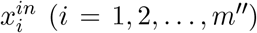 is the intensity associated with the *i^th^* gene in the gene-set. *m’* + *m”* amounts to the total number of genes in the gene-profile. The computation of the U-*s*tatistic requires that the values 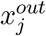 and 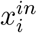 are combined and then rank-transformed. The rank-sum test statistics reported in [11] helps to speed up the computation of the *U*-statistic (then Mann-Whitney-Wilcoxon (MWW) test).

According to [2], the ratio *U/m’m”* is an unbiased estimator of the probability *P*[*X_in_* > *X_out_*], where *X_in_* ~ *F_in_*(*x*) and *X_out_* ~ *F_out_*(*x*). Given a gene-set, the event *X_in_* > *X_out_* says that” *a gene randomly drawn from the gene-set has an intensity greater than the one of a second gene randomly sampled from outside the gene-set*”. We define the Normalized Enrichment Score (NES) as *P*[*X_in_* > *X_out_*] and the associated *p*-values comes from the MWW-test. Assuming that a gene-profile recapitulates the differential expression of treatment samples versus control, a NES close to 1 means association of the gene-set with the treatment. Instead, a NES close to 0 suggests an association with the control group. This interpretation allows us to restate NES as

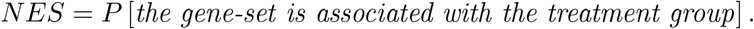

A different way to look at the NES is the odds = NES/(1-NES) that is the imbalance of the probability that the gene-set is associated with the treatment group to the probability that the gene-set has no association with it, or the gene-set is related to the control group.

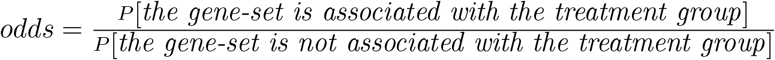

The association with the treatment is as strong as the odds diverges to infinity; it is weak when the odds approaches to zero. In this last case, the association is with the control groups. An odds about 1.0 means no association with either the treatment nor with the control group.

A further transformation of NES is the *logit*2*NES* = log_2_(odds). In this version of the NES, a zero value means no association; a positive value is a measure of the association of the gene-set with the treatment group, while a negative value points at the control.

### 2.2 Enrichments prioritization

It is known that gene-sets with smaller dimension are inclined to get higher NES and lower p-value. The final table of the analysis is usually ranked according to the NES or the p-value, in this way, the attention focuses on marginal significant gene-sets instead of those with larger sizes that could provide a robust understanding of the treatment group. To balance among NES, *p*-value, and the gene-set size, we introduced the recap variable *relevance* (*rel*). Let assume we run a two sided enrichment test so that some gene-sets have *logit*2*NES* > 0, and some others *logit*2*NES* < 0. For the *k^/th^* gene-set, *k’* = 1,2,…, in the collection having *logit*2*NES* > 0, then 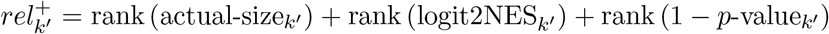, where rank(·) is a function that associates the highest rank with the highest value of its argument, and actual-size is the gene-set size. Similarly, the relevance in the subsets of gene-sets (with index *k*”) such that 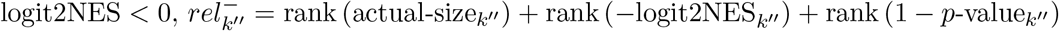. Finally, given the *k^th^* gene-set, its *rel_k_* value is *rel^+^* when logit2NES_*k*_ > 0, and –*rel^−^* when logit2NES_*k*_ < 0. For the ”greater” alternative hypothesis, *rel* ≡ *rel^+^*, and for the “less” hypothesis *rel* ≡ –rel^-^.

### 2.3 Enrichments visualization

We integrate the results with a network-graph of gene-sets. A node represents a significant gene-set. The sizes of the node are proportional to the size of gene-sets, while the intensity of the color is proportional to NES values. The connection between two gene-sets *A* and *B* is instead proportional to their similarity *S*(*A, B*). The similarity is computed as a convex combination of the Jaccard, *δ*_0_(*A, B*) = |*A* ⋂ *B*|/|*A* ∪ *B*|, and the overlap, *δ*_1_(*A, B*) = |*A* ⋂ *B*|/ min (|*A*|, |*B*|), indexes. *S*(*A, B*) = *ϵ* · *δ*_1_(*A,B*) + (1 – *ϵ*) · *δ*_0_(*A,B*), with 0 ≤ *ϵ* ≤ 1. When *ϵ* = 0 we get *S*(*A, B*) ≡ *δ*_0_(*A, B*), while *ϵ* = 1 means *S*(*A, B*) ≡ *δ*_1_(*A, B*).

## 3 Results

To run the analysis, the user needs to load two files: a gene-profile (as a two columns tab-separated text format, the gene-name and the associated value), and one or many collection of gene-sets (in .gmt format). The next steps are: 1) set the significance-level of the enrichments (the user can choose between the *p*-value, and two versions of adjusted *p*-values: Benjamini-Hochberk and Bonferroni), and 2) set the threshold value of the *logit*2*NES*. To require that the probability of association of the gene-set with the treatment group be twice the probability of non association, the *logit*2*NES* threshold must be set to 0.9 (equivalent to *NES* > 0.65, or *odd* > 1.5).

Triggered the computation with the run button, a tabular version of the results is generated (see figure 1). This table respects the constrains given as input, while the full table of the enrichments associated with every gene-set can be downloaded as .csv or .tsv text format. The html version of the table can be downloaded as shown. Both the displayed table, and its .html version, allows the user to re-sort the results according to any of the columns.

**Figure 1:**
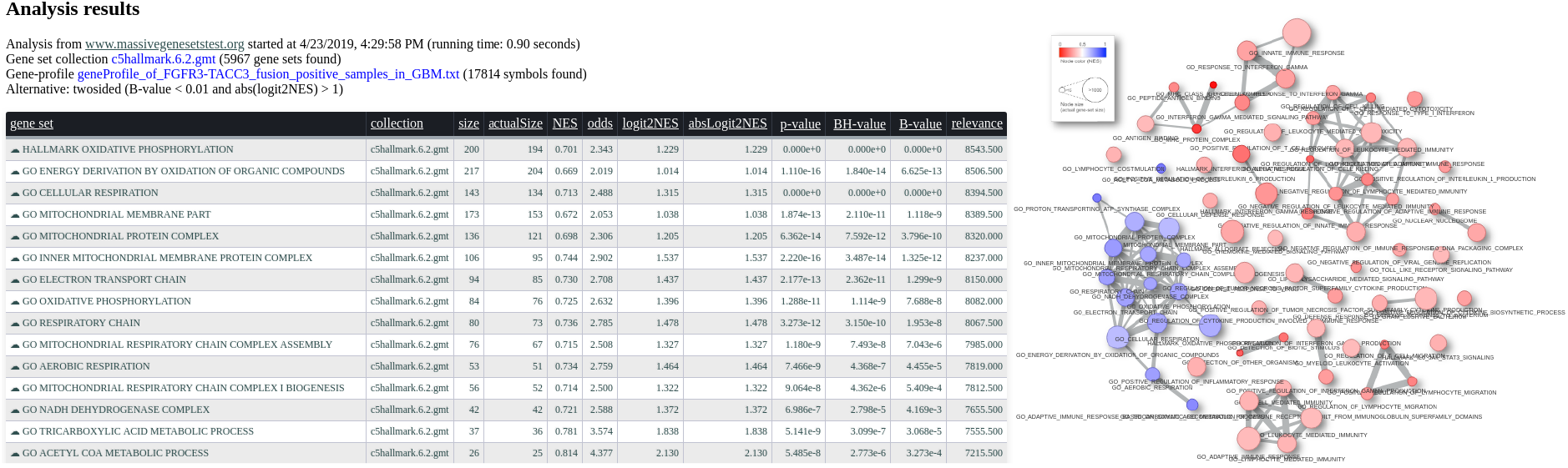
Example of the output of enrichment analysis. On the left, a tabular view of the significant gene-sets; on the right, the same significant gene-sets displayed as a network.

To visualize the network-graph of the current results, the user can click on the network tab. Here, the similarity between any two of the gene-sets in the table is computed and the network of the gene-sets is shown. The user can chose between two similarity measures, the Jaccard or the overlap, or any convex combination of the twos by tuning the parameter e with a slider box. A second slider-box allow to set the threshold value so that a segment joins two nodes when the similarity is above it. As the user operates the two sliders, the network is updated in real time. The plot of the network allows some editing actions and it can be downloaded as .png file.

In figure 1 we present an example of analysis results. We interrogated the gene-profile of the FGFR-TACC3 fusion positive samples in the glioblastoma multiforme study from the TCGA (see [3]) with the C5 and Hallmark collections (MsigDB v.6.1) of gene-sets from the Broad Insitute. On the left, there is a list of the significant gene-sets (alternative = two-sided, B.value < 0.01, and abs(logit2NES) > 1), while the corresponding network is shown on the right. This analysis can be reproduced with the gene-profile provided on the web-site, and gene-sets collections provided by the http://software.broadinstitute.org/gsea/msigdb/index.jspMSigDB.

## Acknowledgements

SMP designed the statistical analysis, and LC engineered the web-site and implemented the statistical functions.

## Availability

massiveGenesetsTest is freely available at http://www.massivegenesetstest.org/

